# What’s next for Indian Ornithology? 101 key research questions

**DOI:** 10.1101/2025.07.25.656497

**Authors:** Devica Ranade, Priti Bangal, Bhoj Kumar Acharya, Peroth Balakrishnan, Sahas Barve, Malyasri Bhattacharya, Aparajita Datta, Mousumi Ghosh-Harihar, Govindan Veeraswami Gopi, Farah Ishtiaq, Soumya Iyengar, Manjari Jain, Girish Jathar, Rajah Jayapal, Panchapakesan Jeganathan, Ashish Jha, Monica Kaushik, Anil Kumar, Raman Kumar, Suresh Kumar, Gopinathan Maheswaran, Shirish S. Manchi, Prachi Mehta, Shomita Mukherjee, PO Nameer, Rohit Naniwadekar, Madhumita Panigrahi, Dhanashree Paranjpe, J Praveen, Suhel Quader, Vivek Ramachandran, Ajai Saxena, Parveen Shaikh, Anusha Shankar, Hilloljyoti Singha, Ankita Sinha, Suryanarayana Subramanya, Kulbhushansingh Suryawanshi, Ashwin Viswanathan

**Affiliations:** Nature Conservation Foundation; Sikkim University; Kerala Forest Research Institute; Archbold Biological Station; Wildlife Institute of India; Tata Institute for Genetics and Society; National Brain Research Centre, Gurugram, Haryana; Indian Institute of Science Education and Research, Mohali; Srushti Conservation Foundation; Salim Ali Centre for Ornithology and Natural History (SACON); Azim Premji University; Zoological Survey of India; Nature Science Initiative; Wildlife Research and Conservation Society, Pune; Kerala Agricultural University; Rupa Rahul Bajaj Centre for Environment and Art; Wildlife Biology and Conservation Program, National Centre for Biological Sciences, Tata Institute of Fundamental Research; IFS (Retd); Bombay Natural History Society; Tata Institute of Fundamental Research Hyderabad; Bodoland University; University of Sheffield; Retired Academic, University of Agricultural Sciences, Bangalore; 1. High Altitude Program, Nature Conservation Foundation, 1311, Amritha, 12th Main, Vijayanagar 1st Stage, Mysore, 570 017 India2. Snow Leopard Trust, Seattle, WA, U.S.A.3. Future Flourishing Program, CIFAR, MaRS Centre, West Tower, 661 University Avenu

## Abstract

India has a rich history of ornithology, and bird research in this field has expanded considerably in recent decades, spurred by a growing number of birdwatchers and ornithologists. Despite this progress, critical gaps remain. This paper highlights key areas where further research is needed and identifies pressing questions that will shape Indian ornithology in the coming years. Drawing from diverse inputs, we present a curated list of 101 research questions spanning across the various disciplines in ornithology – Natural History, Physiology and Disease Ecology, Behaviour, Population and Community Ecology, Habitat Ecology, Macroecology and Biogeography, Population Genetics and Evolution, Applied/Economic Ornithology and Conservation. The list was compiled through a multi-stage process, starting with a public open call for questions, followed by review and curation by a smaller panel of subject specialists. Each question was independently scored by a panel of experts based on three criteria: generality, novelty, and relevance. To account for variation in scoring styles, scores were normalised using Z-scores. The top 101 research questions were then selected based on these standardised scores. Each question is accompanied by an annotation that describes the significance of the question, and highlights opportunities to address it. While our list of research questions highlights significant research priorities, it is not intended to be exhaustive. Rather, it reflects the perspectives of those involved in its curation, who deemed these questions particularly relevant and impactful towards advancing Indian ornithology. We expect these questions to spark new project ideas among students, researchers, and citizen scientists, while guiding funders, managers, and policymakers toward priority research areas.

## Introduction

Ornithology in India has a long history (Ali 1979). Ancient texts contain several observations of birds: for example, the Rig Veda (composed c. 1900–1200 BCE) refers to koels as *anya-vapa*, those raised by others (Friedmann 1965) and there are numerous references to birds in the Sangam literature (roughly from 300–200 BCE to 300 CE) (Varadarajan 1969). Several Mughal emperors, notably Jehangir (years of rule, 1605–1627), were enthusiastic naturalists, compiling natural history notes in their memoirs and commissioning paintings of birds and other wildlife (Ali 1979, Bhat 2020). In the 18th century, after the arrival of European colonists, systematic study of the region’s avifauna led to an expansion in work on taxonomy, nomenclature and natural history, with the first book devoted to Indian birds published in 1862–1864 by TC Jerdon (Pittie 2016). By the time of India’s independence in 1947, much had been published about India’s birds in journals and books, marking a maturation of the formal discipline of ornithology (the academic investigation of topics like evolution, behaviour and physiology with birds as a focus).

The past several decades have seen continued growth of popular and academic interest in birds in India. Just as more people have picked up binoculars and cameras to become birdwatchers and bird photographers (Shyamal 2007, SoIB 2020), so have more scientists been engaged in research on birds. Scientific papers on birds are being published by both amateurs with keen interest and high expertise, as well as by professional researchers. This work is sometimes carried out independently and sometimes under the umbrella of an institution, including universities, government research institutions, and research and conservation organisations in the non-profit sector. For example, the latest State of India’s Birds report was produced as a partnership among researchers from 13 government and non-government organisations, who collectively analyzed data of >70% of the country’s avifauna for conservation prioritization (SoIB 2023). Ornithologists have begun to organise themselves (e.g., the Association of Avian Biologists in India, AABI: www.aabi.in), and directories have been created to connect students with researchers (e.g., <u>ornithology.in/ornithologists-directory</u>).

The long history of work on birds in India, together with the explosive growth over the past few years might suggest that most key questions in Indian ornithology have already been addressed. However, this does not appear to be the case. With a megadiverse avifauna (1,375 species reported at the time of writing (Praveen 2025), most species in India are still poorly known and studied. India contains four biodiversity hotspots and is also an important region for migratory birds that pass across and winter in the country through the Central Asian Flyway (Kumar and Alam 2023). The diversity of habitats and biogeographical contexts provide an opportunity to address many ecological and evolutionary questions. Furthermore, the pace and scale of various human-induced changes necessitate research that supports both conservation theory and practice. Birds in India face several threats ranging from urbanisation and forest degradation to hunting and climate change (Jambhekar et al. 2025). The recent State of India’s Birds report assesses the conservation status of birds in India and summarises what is known about major threats (SoIB 2023).

This combination of circumstances (increasing numbers of researchers, together with the opportunities and requirements for research) prompted an effort to compile a set of key research questions in Indian ornithology. Inspired by previous authors (Sutherland et al. 2006, 2013; Coleman et al. 2019, Cooke et al. 2021), we came together to solicit ideas for research questions from a diverse audience. We then condensed the contributions into a set of 101 research questions. To explain the motivation behind each question, and some opportunities to address it, we have added a brief annotation to each question.

We emphasise that this list is not the only set of questions of importance; merely that the group of people who curated them (i.e., the authors of this paper, most of whom are practising ornithologists) found them to be of particular interest and value.

We offer these questions to underline how much research remains to be done in various fields. We also envision this list to be useful to students and researchers searching for interesting and meaningful topics and questions to choose for their theses, dissertations, or other projects. For this audience, we hope that the annotations will prove particularly useful.

## Methods

The process we followed was to invite questions, to sort and curate them, to prioritise them, and finally to add annotations.

### Inviting questions

In June 2021, an initial group of five individuals (DR, FI, MJ, RJ, SQ) convened to discuss the idea of the project and initiated the process by soliciting research questions from the broader community. Contributions were invited through an online form (see Supplementary Material for details), which was disseminated via email, newsletters, and social media to reach a diverse audience, including nature enthusiasts, students, researchers, and managers. The form encouraged respondents to propose what they considered to be important research questions in Indian ornithology. Submitted questions were required to meet the following criteria:

1. The question should allow for a factual answer, attainable through a realistic research design.
2. The question should address an important gap in knowledge.
3. The question should not be formulated as a general topic area, nor should it be too specific (e.g., to a single species).
4. The question should not likely be answerable with a simple yes or no.

For each response that was submitted, contributors also assigned a subject area they thought was most appropriate from among the following subject areas: Applied/Economic Ornithology, Behaviour, Community Ecology, Conservation, Disease Ecology, Habitat Ecology, Macroecology and Biogeography, Natural History, Physiology, Population Ecology, Systematics and Evolution. Contributors were also asked to indicate their profession or involvement with ornithology: researcher, ecology student, birdwatcher, other nature enthusiast, environmental manager, and others. During analysis, we clubbed birdwatchers and other nature enthusiasts into a single category of ‘Nature Enthusiasts’. A contributor could submit as many responses as they liked.

The form was kept open from 11 September 2021 to 21 November 2021. In all, 782 responses were received from 306 respondents across the 11 predefined subject areas.

At a later stage, these 11 subject areas were collapsed into nine. Population Ecology (9.2% of responses) and Community Ecology (7% of responses) did not receive many responses independently and were merged into the single subject category ‘Population and Community Ecology’. For similar reasons, Disease Ecology (2.7% responses) and Physiology (3.7%) were merged into ‘Physiology and Disease Ecology’. In addition, Systematics and Evolution was renamed ‘Population Genetics and Evolution’ since very few Systematics responses were received. Different audiences contributed a mix of responses across all subject types (Figure 1). The number of responses received from ecology and ornithology researchers/teachers (310 responses), was comparable to those received from nature enthusiasts (325 responses), revealing interest in bird research among non-professionals. The subject area that received the most responses was Conservation (194 responses, c.25% of all responses), and this was true both for responses received from researchers and students (28% of responses this group contributed) and from enthusiasts (23% of responses contributed by enthusiasts).

**Fig 1:**
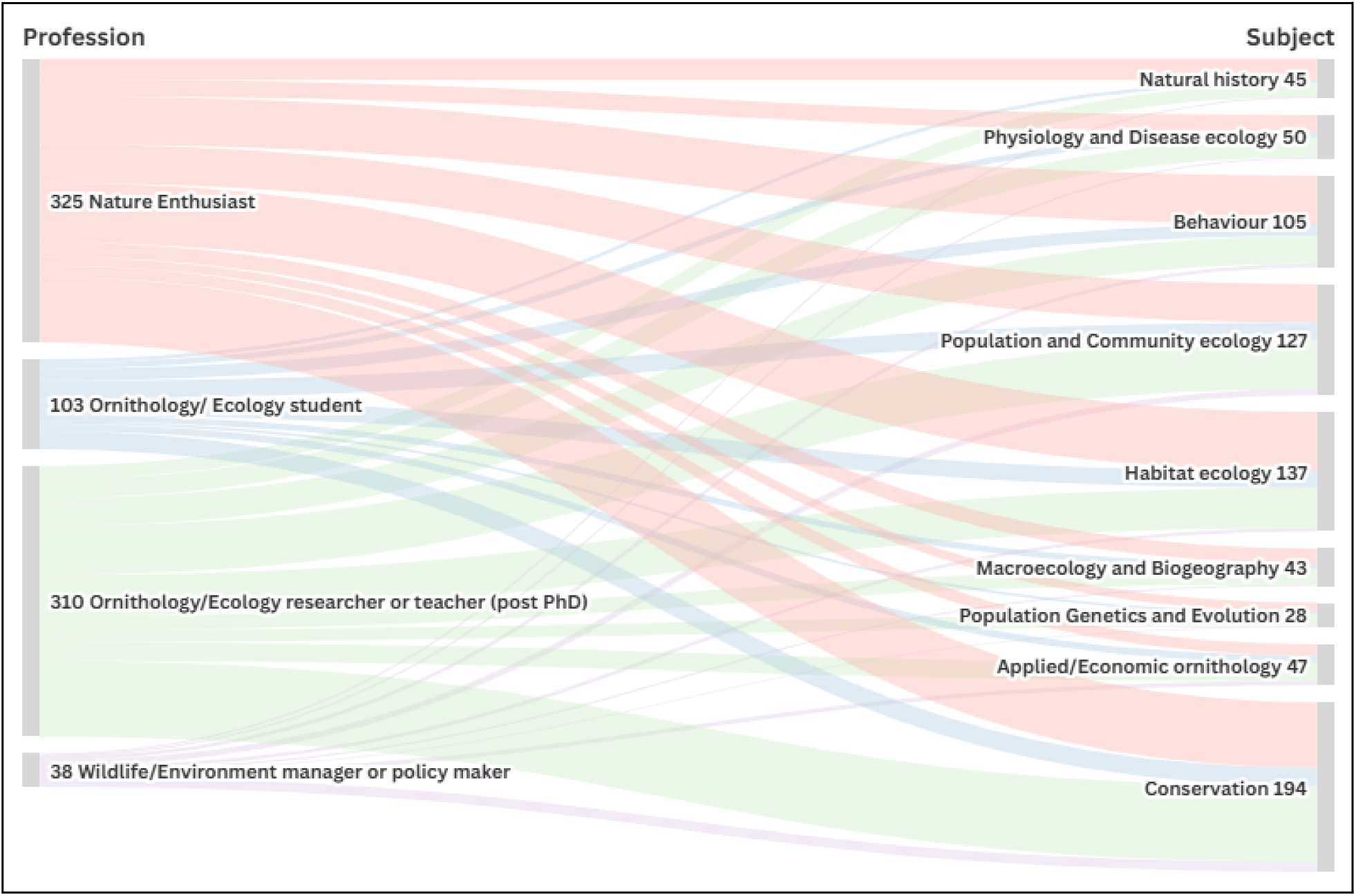
A Sankey plot showing the number of responses by contributors from different backgrounds under the different subject areas. Numbers on the left are the number of responders in each profession category while the number on the right are the number of responses received in each subject category.

### Sorting and curating questions

In May 2022, the initiating set of people invited a larger group to help advance the project. This included researchers, conservationists, and managers, primarily (but not exclusively) selected from the 306 respondents who had contributed responses in Step 1. Finally, a group of 39 individuals was constituted, with a remit to sort, curate and prioritise the contributed responses. Responses in each subject area were examined by a subject working group consisting of between four (Disease Ecology) and eight (Natural History, Systematics and Evolution) members. One person from every subject group was identified as a point of contact and to coordinate group tasks. Two additional working groups were formed, one for workflow and data management, and another to define scoring criteria.

During the course of multiple online meetings, the subject groups examined all 782 contributed responses. They identified and removed responses which clearly did not fit the predefined criteria (195 responses removed). They also identified responses that were similar to each other and could be merged; as well as complex responses that were best split into two (or more) questions. Responses that appeared to have been misclassified by the contributors, and were deemed more appropriate to be placed in a different subject area were moved accordingly (121 responses were moved into a different subject category). In cases where a response appeared to fit into multiple subjects, it was copied into all such subject categories for scrutiny by the individual subject groups (95 responses were placed under more than one subject category). In some cases the subject groups rephrased the responses in their category for better clarity. This process resulted in a longlist of 456 research questions in all.

### Prioritising (scoring) questions

A working group of three members was formed to draft the criteria for scoring questions. Their proposal was discussed by the larger group, and three scoring axes were identified by consensus.

A. **Generality:** How well is a question generalizable to other taxa, and habitats? Does it provide insights about broad-scale patterns and processes?
B. **Novelty:** Has the question been investigated globally or nationally? Does it generate crucial information on understudied groups and/or systems? Does the investigation require innovative methods and approaches like experimental and interdisciplinary work? Is the question related to an emerging avenue of research like climate change, diseases and epidemiology, etc?
C. **Relevance:** How urgent is the need to address the question? Is the question relevant to state, national or international priorities (e.g., state or national policies, Sustainable Development Goals)? How important is addressing the question for local communities, conservation practitioners and policymakers? Is the target species or group of birds of conservation concern? How relevant is the question for the advancement of science (e.g., is it likely to lead to a new area of investigation)?

Members of the subject groups independently evaluated each question in their subject category by providing a rating of 1 (unsatisfactory) to 5 (exemplary) for each of the three criteria: generality, novelty and relevance. The three ratings were then summed to give a single score (between 3 and 15) for each question by a member. All questions within a subject area were evaluated by all members of the relevant working group. The most straightforward way to generate a unique score for each question would be to take a simple average of all the scores that question received. However, some people may be more generous than others on average; and some people may use the entire range of possible scores, which others may not. To account for these possible differences among scorers, the score associated with each person’s evaluation of a question was converted into a Z value by subtracting the mean (of that person’s scores in that subject area) from it, and dividing by the standard deviation (of that person’s scores in that subject area). These Z values measure by how many standard deviations a particular score differs from that scorer’s mean. For example, a Z value of 1.5 for a question would indicate a score that is 1.5 standard deviations higher than the mean of all questions evaluated by that person within that subject area. These Z values were then averaged across all people who scored that question so that all questions within a subject category could then be compared with each other.

On 11 and 12 August 2022, a hybrid (in-person, with an option to join online) meeting of the larger group was held to discuss the scores and chart the course ahead. Twenty-five participants met in person (at National Centre for Biological Sciences (NCBS), Bengaluru) and 6 participants joined online. Participants examined all questions together, and discussed and critiqued the ranks and scores. On day 2, participants split into their respective subject groups and made a list of top questions based on the average Z scores as well as feedback received from the larger group on the previous day. In some cases, questions were further rephrased, merged, or moved between subject groups for improved clarity and coherence. A list of 107 top questions emerged from this, which, after further curation, resulted in a final list of 101 questions.

### Annotating questions

During the hybrid meeting described above, it was decided that subject groups would provide annotations for questions in their subject categories. It was first necessary to formulate guidelines for annotating questions in order to maximise consistency and coherence across subjects and questions. Assuming that those reading the annotations would vary from undergraduate students to experienced researchers, we decided to keep them brief and simply written. Annotations would include a context or background to the question, and a description of its importance (i.e., generality, novelty and/or relevance). Where appropriate, annotations could also hint at opportunities to answer a question (e.g., a suitable taxon, location, technology, etc).

Over the course of subsequent online meetings, subject groups used these guidelines to draft annotations of the questions in their subject categories. Discussions around the annotations often brought more clarity to the understanding and phrasing of the question, and occasionally led to the introduction of additional pertinent questions if the group decided that a key topic had been left out.

After subject groups had annotated their questions, the annotations went through multiple rounds of editing for overall consistency of style and length. During the course of this process, a small number of additional changes were made in terms of merging, rephrasing, and moving questions across subject categories.

## Results

Most questions in our final list are broadly phrased in the following structure: “What pattern of variation in this trait or property (e.g., song, or species diversity) do we see in relation to some external factor (e.g., latitude, or urbanisation).” Sometimes the question is explicitly causal: “What is the effect of this possible cause (e.g., noise pollution, or agrochemicals) on this outcome of interest (e.g., vocalisations, or population change).” In some cases, specific putative causes are named; in other cases, these are left open. Sometimes, specific geographies or environmental conditions (e.g., the Himalaya) are identified as being of particular relevance for the question. On occasion, a more general question is followed by a question that is very similar, but identifies specific conditions of interest. We have deliberately avoided standardizing the questions to position them along a specific-to-general spectrum. Instead, by varying the types and phrasing of the questions, we aim to highlight the richness of potential topics for exploration.

Grouping these questions into different topics is not intended to suggest strict boundaries between subject areas. On the contrary many subject areas blend into each other; and several of the research questions listed here span two or more subject areas. The grouping presented here is for ease of reference, and we have assigned questions to subject areas in a manner we found most fitting, and therefore have occasionally adjusted the classification suggested by the original contributors.

### Natural History

Most in-depth questions in ornithology have their roots in natural history observations, which are basic descriptions of various aspects of the biology of birds, including observational descriptions as well as assessments of various forms of diversity (behavioural, phenological, geographical, etc.). In the past few decades, such natural history studies have gone out of fashion when compared with hypothesis-driven, experimental or technology-dependent research. However, virtually all organismal research is based on a foundation of natural history, hence we devote a section to this theme. We acknowledge that some questions in this section could possibly be placed in others (e.g., Behaviour) and vice versa, because of a general lack of agreement on a precise definition of natural history.

1. What are the patterns and variations in life history traits of India’s birds?

**Annotation:** Understanding fundamental life history traits like clutch size, growth rate of chicks, predation rates on juveniles and adults, lifespan, nesting attempts per year, natal dispersal and more, has been a topic of considerable investigation globally. By and large, species in tropical (= aseasonal temperature) and temperate (= highly seasonal temperature) environments broadly differ in these traits. India spans a large gradient between these extremes, and varies in rainfall seasonality as well. This provides an opportunity to investigate life history traits across these gradients, both among as well as within species, while at the same time filling a crucial knowledge gap in the natural history of Indian birds *(see also question 36)*.

2. How do avian life history traits respond to a gradient of urbanisation?

**Annotation:** Many birds depend on certain features of the habitat (e.g., presence of wetlands, trees with cavities) or host plants for foraging, predator avoidance or reproduction. Such features affect species’ traits such as lifespan, clutch size, growth rates, and more. The alteration of habitats accompanying urbanisation is therefore expected to result in a change in these life history traits as well. Understanding these effects are of interest in their own right, and may also help in the design of sustainable cities that can meet the needs of both humans and biodiversity.

3. How do activity budgets of birds change across land-use gradients?

**Annotation:** A time-activity budget is a quantitative estimate of the proportion of time spent in specific behaviours such as foraging, grooming, resting, vigilance, seeking mates and more. Time-activity budgets vary across species, habitats, and seasons. Different habitat characteristics (often resulting from different human land-use) can significantly alter food availability and habitat features. When food is abundant, birds may spend less time searching for food and extra time can be devoted to other behaviours. During food scarcity, birds could spend more time searching for food, potentially decreasing time available for grooming, social interactions, and vigilance. Similarly, disturbance, predator density, availability of nesting sites are expected to alter time-activity budgets. Because the breakup of how birds allocate their time can affect survival and reproduction, it is important to study how these vary across land-use gradients.

4. How do vocalisations vary geographically within species?

**Annotation:** Song is a highly evolved acoustic signal used to communicate individual identity, territory ownership, and reproductive capabilities. Its structure varies as a result of various evolutionary processes. Documenting variation in birdsong is an important step toward revealing the evolution of reproductive isolation over evolutionary time scales, which may eventually lead to population divergence and even speciation. Geographical variation in song is known in species with disjunct distributions, but is also observed in some species that have a continuous distribution. Widely distributed species of robins, prinias, and whistling-thrushes may be good models for understanding variation in birdsong and its underlying causes.

5. Why do some birds breed solitarily and others in colonies?

**Annotation:** Whether to breed alone or in colonies is an evolved trait that is assumed to result from a tradeoff between costs and benefits. Some bird families are almost entirely social in breeding (e.g., weavers), while other families (e.g., storks) have both solitary and colonial breeding species. There may be intraspecific variation as well, with some individuals breeding solitarily and others in colonies (e.g., bee-eaters). Many factors can favour colonial nesting – predation pressures, coordinated access to food supplies, or lack of adequate nesting sites.

However, this is at the cost of competition, sharing of scarce resources, and transmission of disease. Research is needed to identify the costs and benefits of colonial versus solitary nesting in Indian birds, as well as how they vary across different species and environmental contexts *(see also question 21)*.

6. What influences patterns of extra-pair parentage?

**Annotation:** Studies on avian mating systems have shown that true monogamy is rare. Even socially monogamous species are known to be sexually promiscuous, with high rates of extra-pair paternity. What drives the evolution of such a mixed reproductive strategy, with the social and true mating system being different? This is an area of active research but with virtually no inputs from Indian model systems. There is an opportunity to understand how life history traits (parental investment, clutch size, longevity, etc) might act in combination with other factors (like population density) to explain the prevalence of and variation in extra-pair paternity.

7. What factors influence breeding success of birds in the Himalaya?

**Annotation:** Several factors influence breeding success in birds, including the availability of appropriate food and nesting sites, predation pressure, and prevalence of parasites and disease. In the Himalaya, unfavourable weather conditions (such as low temperature, snow and scarcity of food), may increase the challenges and impose additional costs for successful breeding. How do these environmental factors act directly and indirectly to affect the seasonal timing of breeding of birds, and consequent breeding success?

Physiology and Disease Ecology (12 Questions)

Most of the research themes listed in this paper are at the whole-organism level or above; however there is still much to be learnt about underpinning mechanisms, including physiology and neurobiology. In addition, avian disease and its ecological influences is a topic of major contemporary research, both in its own right, as well as in its connections with human wellbeing. These themes are highly under-studied in the Indian context, and many fascinating research opportunities exist.

8. What physiological and behavioural adaptations allow birds to persist in different climatic zones?

**Annotation:** Birds can adapt to different climatic conditions (e.g. hot deserts vs. tropical rainforests), in very different ways. Some of their physiological parameters can shift; for instance their basal metabolic rates or evaporative cooling rates might have to be adjusted given the temperature differences. In parallel, or separately, they might make behavioural adjustments, for instance by spending more or less time in the shade. One way to study such a question could be to assess closely related species that live in very different environments, or to study a migratory species that spends some time in different environments.

9. How does diet influence the gut microbiome of birds?

**Annotation:** Several studies have revealed the importance of microbes in the development and immunity of their host species. The composition of gut bacterial communities is likely to be influenced by factors such as habitat type, species associations and interactions, gut anatomy, host genetics and diet. Indian birds offer excellent model systems to study this given the variety of habitats, social systems and diversity of avian communities present in Indian landscapes.

Such studies are largely missing and can help understand the effect of provisioning on birds, in addition to the evolution of host-microbe associations in Indian avian systems. The gut microbiome can also be studied for the same species in different habitats - for instance across wintering and breeding grounds in migratory species.

10. How are the energy budgets of birds expected to change with climate change?

**Annotation:** Birds spend energy on resting, moving, sleeping, foraging, migrating, and so on. As aspects of their environment (temperature, rainfall, food availability, etc.) change, they have to adapt what they spend their time and energy on (i.e., activity and energy budgets) to meet their daily energetic needs. For instance, if food availability decreases in their environment, they can either spend more time being active and searching for food, or spend more time resting to conserve energy. With climate change, food availability, temperature, and rainfall patterns can all shift, causing shifts in energy and activity budgets; and this is an area that requires research attention.

11. How do anthropogenic changes impact the body condition and stress physiology of birds?

**Annotation:** Human-made environmental changes such as changes to land use patterns, or rural-urban gradients, can impact many aspects of bird physiology. These aspects can be indicators of body condition, such as parasite load, feather quality, immune function, etc., or indicators of stress, such as corticosterone levels. These changes can operate through a number of factors, including nest site availability, food quality and diversity, pollution (air, water, light, sound, etc.). For instance, how are body condition and stress affected across rural-urban gradients in a common bird like Jungle Babbler or Common Myna?

12. What is the neurobiological basis of song learning, vocalisation and song recognition in birds?

**Annotation:** Songbirds are known to learn their songs during a sensitive period during early development from adult conspecific tutors. Specialised neural circuits, called the song control system, are important for song learning, vocalisation and perception in commonly studied songbirds such as zebra finches *Taeniopygia* and canaries *Serinus canaria*. However, fewer studies exist on brain-behaviour interactions for singing in various species of Indian songbirds. Besides field studies to record songs sung in different contexts, investigating the structure and functions of the song control circuitry can be undertaken to elucidate the neurobiological basis of singing and vocalisation in Indian songbirds.

13. How do micronutrients affect avian reproductive success?

**Annotation:** Micronutrients, including vitamins and minerals, constitute a small fraction of the diet, but are extremely important for the overall health of birds, especially in reproductive success. Deficiencies in minerals lead to reduced fertility of both sexes and also decreases in the chances of their eggs hatching and in the survival of the progeny. Calcium, Phosphorus, Sodium and Potassium are known to be vital for the viability of breeding birds and for the formation of egg shells. Zinc, Copper, Manganese and Selenium can indirectly affect reproductive success by affecting feeding and the humoral immune system of birds. Whereas the levels of micronutrients have been optimised mainly for poultry, such data are scarce for wild birds and this needs to be studied further.

14. How does light and sound pollution impact bird physiology and sensory ecology?

**Annotation:** Artificial light at night (ALAN) and noise pollution alter the sensory environment of animals and are tightly associated with urbanisation. Mounting evidence points towards their negative impacts on wildlife. ALAN can result in disorientation, exhaustion or even mortality from collisions in migratory birds. In songbirds high noise levels can result in changes in sound production or perception, including hearing loss. Both noise and light pollution may also affect the circadian rhythm and stress levels in birds. India is undergoing rapid urbanisation, making it imperative to investigate the effects of light and sound pollution on birds, and ways to mitigate them *(see also questions 33 and 49)*.

15. What are the physiological impacts of chemical pollutants on the health of birds at different trophic levels?

**Annotation:** Environmental contamination by agricultural chemicals and industrial waste disposal results in adverse effects on the reproductive success of exposed birds. The diversity of pollutants results in physiological effects at several levels, including direct effects on breeding adults as well as developmental effects on embryos. The range of chemical effects on adult birds includes acute mortality, sublethal stress, reduced fertility, eggshell thinning, and impaired incubation and chick rearing behaviours. Furthermore, chemical pollutants affect birds with different diets from different habitats: diet plays a role in exposure to various chemical pollutants and accumulation across the food chain (e.g., carnivores to scavengers; even low amounts of DDT in water could reach a high concentration in fish-eating birds through biomagnification).

The summary presented here is from global research; work in India lags far behind.

16. What are the spatial and temporal patterns of bird flu in wild species, and how can this be effectively monitored?

**Annotation:** Avian influenza (bird flu) type A is a highly contagious viral disease caused by multiple strains of influenza virus. Birds can be infected with low pathogenic avian influenza virus (LPAI: H1-H16) which is common in wild birds. In poultry, it is a mild disease, with symptoms including mild breathing problems and reduced egg production. High pathogenic avian influenza virus (HPAI: only H5 and H7 subtypes) causes outbreaks and associated mortality in poultry and wild birds. HPAI, subtype H5N1, constitutes one of the world’s most serious health and economic concerns given the catastrophic impact of epizootics on the poultry industry, the high mortality attending spill over in humans, and its potential as a source subtype for a future pandemic. Nevertheless, we still lack an adequate understanding of HPAI H5N1 epidemiology and infection ecology among wild birds.

17. What is the role of birds as carriers of antibiotic resistant bacteria?

**Annotation:** As in all vertebrates, trillions of microbes live in the guts of birds and aid in nutrition, immunity, and gut development of their hosts. Furthermore, several studies demonstrate that migratory birds both carry and acquire antimicrobial resistant bacteria from contaminated environments near areas with high poultry or human densities and transmit the bacteria along migration flyways. There is no information on microbial composition and associated antimicrobial genes within the guts of migratory birds (e.g., ducks and geese); given the potential for transmissibility over long distances, this is a crucial area for research.

18. How does disease in hyperabundant species affect humans and wild birds?

**Annotation:** Places with dense populations of hyperabundant species like poultry farms, pet markets, or urban pigeon nesting sites—can serve as hotspots for infectious agents, such as bacteria, fungi, protozoa, and viruses. While most avian diseases do not pose a serious threat to people and wild birds, the severity of the disease is governed by the virulence of the infectious agent, host immunity, and the route of disease transmission. Currently, we lack such data on Indian context and impact of such spillover (if any) from such hyperabundant aggregations to humans and wild birds.

19. What is the extent and function of olfaction in Indian birds?

**Annotation:** Until recently ornithologists believed that birds are largely unable to detect and use olfactory information. However, recent anatomical, neuroanatomical, physiological and behavioural studies have demonstrated the complexity of avian olfactory structures and the functional role of olfaction in orientation, foraging, recognition of familiar odours, predator detection, nest recognition, control of reproductive behaviour, and more. However, such studies are completely lacking in India. Both field and experimental studies on the sense of olfaction, including morphology and neuroanatomy; neural pathways mediating olfactory stimuli and functional role of olfaction in various behavioural contexts will help to advance the current understanding of this field.

### Behaviour (15 Questions)

Worldwide, birds have been a favourite subject for the study of both the mechanisms as well as the adaptiveness of behaviour. Their ubiquity, variation in plumage and courtship behaviour, as well as diversity in habitat, feeding and social behaviour makes them excellent subjects to address a number of questions. Behavioural research on birds has a long history; yet has still only scratched the surface of what is possible.

20. How do birds learn to avoid aposematic prey?

**Annotation:** Several insects are known to sequester plant toxins in their bodies. When birds eat these insects, they fall ill and learn to avoid these insects subsequently. Some birds also just avoid eating novel prey (a phenomenon known as neophobia) and brightly coloured prey. The mechanisms of avoiding toxic insects (whether this behaviour is learnt by copying, by experience, or is instinctive), has been a topic of interest for several decades in behavioural ecology. The study of prey colours and avoidance behaviour of predators serves the broader purpose of understanding the coevolution in predator-prey systems, a phenomenon that remains to be studied in Indian birds.

21. What are the ecological drivers of the evolution of cooperative breeding in Indian birds?

**Annotation:** Cooperative breeders live in stable social groups where some individuals (helpers) forgo their own reproduction to care for offspring of others (breeders). This fascinating reproductive strategy occurs in fewer than 10% of the world’s birds. India is home to many cooperative bird species that occupy diverse habitats. Yet, long-term studies on cooperatively breeding Indian birds are largely lacking. Studying their behaviour under different ecological conditions will allow a dissection of the roles of various social and ecological factors in the evolution of cooperative breeding *(see also question 5)*.

22. What are the ecological and evolutionary underpinnings of brood parasitism in Indian birds?

**Annotation:** Brood parasitism is a key reproductive strategy for some bird species and has been a subject of several studies, but largely in temperate regions. Brood parasitism involves an array of ecological and evolutionary interrelationships including the coevolution of host defences and parasite counter-defences at different stages of the breeding cycle. Hosts typically incur a fitness cost, with consequent implications for the population trends of the species involved.

However, there is dearth of information on brood parasitism of Indian birds, including basic host-parasite associations. In this context, systematic studies on the ecological and evolutionary relationships of brood parasitism is a key issue to be addressed.

23. How does song complexity and function vary between the sexes in Indian species in which females sing?

**Annotation:** Given recent evidence that female song is ancestral and widespread in songbirds, we must reevaluate the assumption that bird song functions primarily in female choice or male-male competition. This calls for a large-scale evaluation of the function and complexity of female bird song through comparative studies across species, and for an examination of the differences in structure and function of song between sexes within a species. Indian birds provide an opportunity to understand the factors that may have driven the maintenance or loss of song in female birds *(see also question 74)*.

24. How does mate choice shape the evolution of courtship signals in birds?

**Annotation:** A large number of studies on sexual selection in avian species have focussed on identifying physical characteristics (secondary sexual characters) that play an important role in intrasexual competition or mate choice. Such characters and behaviours associated with courtship or courtship displays are important dynamic characters that can be targets of sexual selection via mate choice. These inherently complex dynamic signals combine different modalities such as visual (e.g., movements, colours) and aural (e.g., song). In the context of mate choice, these modalities can be complementary (each adding information to the other) or redundant (each substituting for the other). Studies on the detailed role of each signal (e.g., spectral reflectance of plumage; spectro-temporal features and complexity of song) in mate choice will be fascinating. In addition, how these combine to influence the evolution of complex multimodal signalling needs to be investigated in many more avian species.

25. How do personality traits influence mate choice?

**Annotation:** Just as humans do, animals can display personality traits – suites of consistent behaviour that range across individuals along a spectrum of boldness, activity, sociability, tendency towards exploratory or aggressive behaviour, and so on. These suites of traits are consistently observable across time and different contexts, and give a unique ‘personality’ to individuals. Whether these personality traits play a role in mate choice remains to be seen. Although it might be difficult to experimentally manipulate personality traits, choice experiments between different personality types may provide valuable insights into how behavioural differences can contribute to mate choice in birds.

26. How does non-consumptive predation risk affect behavioural responses of birds?

**Annotation:** Predation risk is known to affect how prey use habitats in the landscape they live in. However, in addition to the actual risk of being captured and eaten, prey movement and behaviour is also likely to be influenced by the perceived predation risk. This is also known as non-consumptive predation risk. This is a relatively new field of study and understanding how fear drives habitat use and behaviour of birds is a field of research that has generated interest among biologists over the last few years. Possible model systems include pigeon behaviour in response to risk of predation by falcons, or behaviour of small passerines in response to hawks.

27. How do migratory birds learn migratory routes?

**Annotation:** Birds migrate to escape extreme climate and to track food resources. But how do they know where to go? Previous research suggests that both genetic instincts and social learning play a role. Indian birds exhibit a diversity of migratory patterns, from seasonal and elevational migration to long-distance latitudinal migration. Using genomics and stable isotopes to connect populations across seasons is one step to understand the evolution of migration.

Another is to use behavioural experiments on individuals to understand the social, hormonal and climatic cues birds use to initiate or end migration.

28. What are the patterns of site fidelity among migratory birds?

**Annotation:** Over 300 species of birds migrate to, or through, India. In some species, individuals are known to return to the same site (even the same tree) in successive years. But site fidelity to patches may vary within and across species. Widespread and long-term bird ringing programs are important to ascertain how common site fidelity is across all the various taxonomic families that have migratory species, in both breeding and wintering sites. Population level and individual level studies are needed to understand how and why site fidelity to patches changes from year to year. Such studies may be especially important for species of conservation significance.

29. To what extent do migratory species compete with resident birds when they overlap in space and time?

**Annotation:** Migratory birds share space and other resources with resident species when they travel to their migratory grounds. It is likely that resident and migratory birds partition resources (use different resources like different foraging habits, different feeding heights) to avoid competition. However, the extent and consequences of competition between migratory and resident birds has not been studied.

30. Which biotic and abiotic factors influence arrival time and duration of stay in migratory birds?

**Annotation:** Birds are known to migrate to avoid extreme temperatures and food scarcity in harsh seasons. Both biotic (e.g., physiological) and abiotic (e.g., physical) factors are known to influence migratory behaviour. Climate and anthropogenic changes are likely to alter the conditions that are suitable for migratory birds to arrive in and survive at their migratory grounds.

The environmental conditions at migratory grounds are therefore crucial in order to ensure the survival of migratory birds. The Indian subcontinent is home to about 300 migratory species, some of whose global populations (e.g., Rosy Starling *Pastor roseus*, Red-breasted Flycatcher *Ficedula parva*) is hosted here during the winter months. It is therefore important to understand what factors influence when birds arrive at their migratory grounds and how long they stay.

31. How does climate change mediate bird phenology?

**Annotation:** The timings of seasonal cyclical events such as moulting, migration, and behavioural changes associated with breeding are known to be influenced by changes in temperature, day length, rainfall patterns and other immediate environmental factors. However, whether and how long-term changes in the local climatic conditions will shift the timings of these phenological phases remains to be seen. Although some information is available about this from temperate parts of the globe, long-term data sets from the tropical and subtropical regions of India are needed to shed light on changes associated with phenological events such as onset of breeding, schedules of migration and even suitability of wintering/breeding grounds under changing climate patterns. One phenomenon of interest is elevational migration (e.g., in the Himalaya), which is expected to be particularly sensitive to climate change over short spans of time.

32. What kind of behavioural changes allow birds to respond to changing landscapes?

**Annotation:** The sustenance and survival of any species depends on successful foraging and breeding in its environment. A problem arises when the landscape changes rapidly as a result of anthropogenic activities or natural disasters. In such cases, individuals may either shift to a more favourable landscape or habitat nearby, or change their behaviour as the situation demands (i.e., by showing behavioural plasticity). To understand what enables the persistence of avifauna in a changing landscape, it is necessary to study a species’ behavioural changes in foraging (i.e., food selection) and breeding (i.e., nest site selection, nesting material, and nest architecture). Suitable model species for such studies are common and widespread species like bulbuls, mynas, swallows, and munias.

33. What is the impact of artificial light on the foraging and breeding behaviour of birds in urban landscapes?

**Annotation:** Artificial light at night is an increasingly important feature of urban habitats. Such light may change activity patterns of both diurnal and nocturnal birds. As cities expand, it is important to study how night lights affect bird behaviour. For example, how do diurnal birds change their activity budgets in response to night lights, and how does that in turn change their foraging and reproductive success? In addition, the costs and benefits of artificial night lights to nocturnal birds also remain to be understood *(see also questions 14 and 49)*.

34. How do birds communicate in noisy acoustic environments?

**Annotation:** When animals communicate using sound in noisy environments (background noise from other calling animals, wind, water or traffic), their sounds overlap with those from the background, resulting in masking. Masking is a serious problem because it makes it harder for receivers to extract meaningful sounds from irrelevant noise. Birds may respond to masking using a variety of strategies, including increasing signal amplitude (i.e., vocalise loudly) and frequency, using more pure tones, and shifting song to times of less noise, such as to the night. Understanding how birds overcome masking will help understand how acoustic niches are partitioned and may generate ideas for signal discrimination for human applications. It is also important to understand how bird vocalisations are affected by anthropogenic noise, for example in urban environments.

### Population and Community Ecology (13 Questions)

The causes and consequences of the dynamics of populations is a fundamental area of ornithological investigation. How are changes in population related to food, habitat, predation, and so on? Can we understand the details of these processes by looking at underlying demographic factors like rates of survival, reproduction, immigration and emigration? How are these factors affected by the mix of species that birds are surrounded by, and interact with?

What are the outcomes of these interactions in terms of the functional role that birds play, and what kinds of species can coexist? These population and community ecology questions are often ideally answered through longitudinal (long term) studies, but much can be learnt from cross-sectional (e.g., space-for-time) studies as well.

35. What are the effects of climate change on the demography of birds?

**Annotation:** The impact of climate change on demography of the birds is an understudied subject. Anecdotes on the failure of nesting of species such as Swamp Francolin *Ortygornis gularis*, Black-breasted Parrotbill *Paradoxornis flavirostris* and Marsh Babbler *Pellorneum palustre* due to untimely floods in the river Brahmaputra are attributed to climate change. Similarly, changes in the phenology of Himalayan plants and their impact on nectar-dependent sunbird species are reported by local communities in the higher Himalaya. The duration and extent of monsoon rains are likely to affect several bird species including weavers and munias. However, careful long-term research to understand the effect of climate change on demographic processes (reproduction, survival) in Indian birds is scarce.

36. How does survival and reproductive success vary across different environmental gradients?

**Annotation:** Widespread species typically occur across a range of environmental conditions, including habitat type, habitat quality, elevation, temperature, and rainfall. A key question is how demographic parameters (e.g., survival and reproductive success) vary across these gradients. Understanding the role of abiotic (e.g., precipitation) and biotic (e.g., food, predation) factors in influencing demography will allow better predictions of population change expected under different scenarios of climate and land-use change *(see also question 1)*.

37. What are the drivers and ecological factors that hasten local extinctions of species?

**Annotation:** Local extinction is a process where a species vanishes from a particular geographic area, while persisting in other sites within its distribution. Various drivers may cause the local extinction of a species, including habitat change, climate change, overexploitation, and natural disasters. For example, the Great Indian Bustard *Ardeotis nigriceps*, earlier found in many pockets of Maharashtra, has been driven to local extinction in most of these areas due to habitat change. Causes may differ between species and across sites, and hence understanding the drivers behind local extinctions is essential for finding ways to sustain populations of species.

38. What is the population status of and threats to nocturnal birds in India?

**Annotation:** India harbours more than 40 species of nocturnal birds, including owls (Tytonidae and Strigidae), nightjars (Caprimulgidae), and frogmouths (Podargidae). Their cryptic nature means that nocturnal species are poorly studied in comparison with diurnal birds. Given rapid changes seen among other species, the population status and threat to nocturnal birds needs urgent investigation. What is the status of nocturnal birds, what factors govern distribution and abundance, and what are the impacts of habitat transformation and cultural beliefs (*see question 94*)? These are vital questions to be answered for nocturnal birds.

39. What is the impact of changing insect populations on insectivorous birds?

**Annotation:** A large fraction of bird species depend on insects and other invertebrates (like spiders) as prey. Many other birds that are usually categorised as nectarivorous, omnivorous, or frugivorous might also consume large numbers of insects. Insects are generally in global decline, largely due to agricultural intensification and increase in agrochemical inputs, particularly insecticides. In parallel, insectivorous birds in India are declining on average. Is there a causal connection between the two patterns? What other factors might dampen or strengthen such a relationship? These are important areas of research.

40. How do anthropogenic influences in the non-breeding grounds affect populations of migratory birds?

**Annotation:** Non-breeding grounds play an important role in the annual migration cycle of migratory birds of all kinds. Many migratory species show strong site fidelity towards their non-breeding locations. Any changes or alterations at these locations are likely to affect their feeding behaviour, timing of migration, refuelling abilities and other aspects that are crucial to survival and successful migration. What human-caused pressures are happening in non-breeding grounds, and how do they affect migratory species?

41. What processes influence bird assemblages across environmental gradients?

**Annotation:** At a community level, biotic processes like competition and predation are thought to structure bird communities. At larger scales, climate and geography influence the broad assemblage of species in a region. While the role of biotic processes and of environmental factors are typically studied separately, rapid global land use and climate change mean that there is a need to examine both sets of interactions at a large spatial scale, and across multiple locations.

42. How does land-use change affect bird communities and populations?

**Annotation:** Land-use change is one of the major drivers of biodiversity loss. It can involve largescale change in habitats as well as fragmentation. It is critical to understand the impacts of land-use change on different habitats (e.g. forests, open natural ecosystems) and bird guilds (e.g. feeding, nesting, habitat). Which are the most harmful changes, and which are relatively benign? What are its effects on habitat specialist/sensitive species, and how can these effects be mitigated? These questions are particularly urgent to answer in habitats that are presently undergoing rapid change and fragmentation—for example grasslands.

43. How can urbanisation be planned to facilitate the maintenance of bird diversity?

**Annotation:** Urbanisation results in increased human infrastructure and modified environmental conditions (e.g., light and noise pollution). Natural habitats are typically reduced, fragmented, and highly manicured, becoming unsuitable for many bird species, particularly habitat specialists. Given the high rates of urbanisation, it is critical to determine how urbanisation can be planned to maintain taxonomic and functional diversity of birds.

44. How do infrastructure projects affect bird diversity and populations?

**Annotation:** Infrastructure projects (e.g., power lines, windmills, dams, railway lines, roads, canals) in urban and rural settings can permanently alter habitats and movement patterns of birds, with consequences for bird population and species diversity. Given the rapid increase in developmental projects it is critical to document their impacts on birds, to assess suitable alternatives or mitigation measures, and to monitor the efficacy of their implementation.

45. How does provisioning affect bird communities?

**Annotation:** How are bird communities affected by deliberate provisioning as well as provisioning that is unintentional (e.g., domestic waste)? Provisioning of food (through bird feeders, garbage, and deliberate feeding) and nesting sites (through nest boxes and building structures) is increasing with urbanization and can alter bird behaviour and community composition. While some species benefit from these resources, others may be negatively affected by contaminants, pathogens, or changes in competition dynamics. Landfills and garbage dumps provide abundant food for certain bird groups, sometimes even attracting threatened species (e.g., Greater Adjutants in Guwahati, raptors at Deonar Abattoir in Mumbai). However, such sites may also expose birds to pollutants and diseases. What types of species benefit from deliberate and from inadvertent provisioning, and what could be its detrimental effects?

46. How do exotic and invasive plants affect birds?

**Annotation:** Plant species are introduced intentionally or inadvertently in different habitats (e.g., forest plantations, ornamental/avenue trees in urban areas). Some of these species can become invasive. As exotic plants invade new habitats, they might alter habitat structure and resource (fruits, flowers/nectar, nesting) availability for native bird communities. What are the impacts of such plants on the behaviour, demography, population dynamics, and community traits of birds? *(see also question 47)*

47. What are the consequences of mutualistic interactions between birds and invasive plants on ecological communities?

**Annotation:** Negative interactions between invasive plant species and bird communities have received some research attention. However, in some situations, invasive plants and native birds may benefit from each other. For example, there appears to be a mutualistic relationship between Lantana and many frugivorous birds, which might lead to the ironic outcome of native species facilitating the spread of invasives, thereby altering habitat structure and composition, and changing resource availability for other native species. How do these effects occur, under what circumstances, and with what outcomes? *(see also question 46)*

### Habitat Ecology (12 Questions)

Birds are creatures of habitat. Which species are able to persist, and how they go about their daily lives, is determined to a large extent by habitat. Hence, the relationship between birds and their habitat, as well as various consequences of habitat alteration, are important to investigate. While in the past considerable attention has been paid to terrestrial habitats, especially forests, the importance of other habitats, including grasslands and wetlands, is becoming increasingly clear.

48. How rapidly are different bird habitats changing across India?

**Annotation:** In the last decade, it has become evident that the degradation and loss of habitat are among the key reasons for global bird declines. What is the rate of degradation and loss of key bird habitats across India at different spatial scales? The answer to this question is needed as input into strategies for the preservation of remnant habitats as well as for efforts to reconcile conservation goals with other human needs (such as agriculture, housing, and resource extraction).

49. How does light and noise pollution affect habitat suitability for birds?

**Annotation:** Birds in urban areas are exposed to elevated levels and long spells of light and noise. Noise pollution can greatly affect the suitability of seemingly good habitat by hampering a bird’s ability to detect and respond to acoustic signals. Likewise, light pollution can disrupt the natural patterns of light and dark experienced by birds. Longer than normal exposure to light might also affect habitat by altering seed germination, invertebrate availability, and phenological patterns of plants and invertebrates, thereby indirectly affecting birds. Research is needed to investigate the independent as well as combined effects of noise and light pollution on habitat suitability for birds in urban areas *(see also question 14 and 33)*.

50. How do sea level rise and extreme weather events impact pelagic and shorebird habitats?

**Annotation:** The habitats of pelagic birds are understudied in India. Breeding colonies on Pitti Island (in the Lakshadweep archipelago) and elsewhere may be under threat from sea level rise and extreme weather events. Along the mainland coastline, crucial wintering habitats for migratory shorebirds may similarly be under threat from erosion and habitat alteration accelerated by climate change. How these threats affect breeding numbers and breeding success of birds is not understood, nor are there projections of future scenarios.

51. How are changes to waterscapes of India affecting birds?

**Annotation:** Wetlands are home to a large number of bird species and provide many useful functions for birds and humans alike. Birds use wetlands to breed, feed, roost, and as stopover sites during migration. Humans rely on wetlands for a number of resources, including water and fish. Changing climate patterns and increasing human needs are altering key features of wetlands, such as the level of water and seasonality of resources. What precisely are these changes and how do they affect the diversity, population, breeding seasonality, and migration patterns in birds? These questions need to be addressed separately in different wetland types and for different bird groups *(see also question 52)*.

52. What is the impact of altered river hydromorphology and riparian landuse-landcover change on habitat availability and use by riverine birds?

**Annotation:** Some birds have specialised to feed on aquatic, terrestrial, and semi-aquatic prey found in and near running waters, and to nest in riparian areas. River flow characteristics and land-use of riparian areas are key habitat elements regulating food availability and nesting conditions for riverine birds. Dams and impoundments disrupt the natural flow of rivers, and other human activities modify riparian vegetation and natural riverine bank features, and drown natural islands. Some of these impacts are small-scale and immediate while others are large-scale and long-term. What are these modifications that alter natural riverine habitat structure, and how do these affect habitat selection and population densities of riverine birds? *(see also question 51)*

53. How do changes in habitat and climate at stopover sites alter food availability and survival of migratory birds?

**Annotation:** Habitat structure and complexity determines the availability of resources, including food, and thus influence bird assemblages. Migratory birds require high energy during flight, and many halt at key stopover sites for feeding. Such sites need to be identified, and research carried out to understand how changes in climate and habitat at these sites affect the availability of food and other resources, which have downstream effects on the survival of these species.

54. How do anthropogenic activities affect the birds of open natural ecosystems?

**Annotation:** Open Natural Ecosystems or ONEs (such as deserts, grasslands and shrublands) are understudied in India. They are historically neglected in Government policy, often labelled as ‘wastelands’, and earmarked to be made more ‘productive’. As a result, these are among the most heavily modified and therefore threatened ecosystems in India. Studies are needed to understand how birds of ONEs are affected by human activities, including grazing livestock, fire, infrastructure like windmills and powerlines, and land conversion and fragmentation.

55. How does fire affect birds across different habitats?

**Annotation:** Historically, fire has been part of many natural ecosystems, especially of dry deciduous forests and grasslands. However, increasing warming and fuel load due to invasive plant species (e.g., *Lantana camara*, *Pinus roxburghii*) is resulting in higher frequency and intensity of fires. Additionally, fire is also used as a management tool in both forest and grassland ecosystems. What is the effect of both managed fires and wildfires on birds?

Conversely, what is the effect of suppression of fire, a widespread management goal? The answers to these questions are likely to vary across ecosystems, and this calls for a research agenda that is implemented at a large scale.

56. How do urban structures affect resident and migratory birds?

**Annotation:** Urbanisation alters natural ecosystems by increasing the proportion of built areas together with heightened human population and accompanying energy and resource consumption. Built structures modify food and nesting resources for birds, and alter microclimatic conditions. They also pose a potential threat as obstacles to movement and sources of collision and mortality of birds. Research is needed to understand the direct and indirect effects of urban structures on birds belonging to different functional groups, to reduce potential negative impacts.

57. How does the type and intensity of farming affect birds?

**Annotation:** Agricultural practices encompass various types of farming at different intensities. Agricultural intensification is a multi-faceted process for achieving increased yields (agricultural production per unit area). It usually involves increase in agricultural inputs and changes in farming practices that lead to conversion of natural habitat to cropland, fragmentation of natural habitat, changes in crop types and cycles, reduction in the extent and duration of fallows, and reduction in habitat heterogeneity at multiple spatial scales. While considerable research has been done globally, and a few Indian studies exist as well, the extent and consequences of agriculture require much more research in India. These could address the impacts of intensification of farming on bird diversity (including taxonomic, functional and phylogenetic diversity) at different spatial and temporal scales, and an exploration of different scenarios under which agricultural production and bird conservation can co-exist.

58. How are birds affected by changing irrigation patterns in arid and semi-arid habitats?

**Annotation:** Changes in irrigation influence hydrology and land-use in arid and semi-arid regions through increases of land area under agriculture, alteration of water levels, afforestation along the irrigation channel network, introduction of invasive plant species, and development of seepage wetlands. How do these changes impact the composition, richness, and abundances of birds at different spatial scales? How do habitat specialists, such as certain farmland birds, birds of arid or semi-arid regions (e.g. bustards), wetland birds and also ‘common’ species, respond to changes driven by irrigation? Although some work has been done to address these questions, the immense scale of the transformation calls for further research across a variety of landscapes.

59. How is biotic homogenisation in urban areas affected by the kinds of life history traits that are selected for in cities?

**Annotation:** Recent research in urban areas has shown that cities around the world are occupied by species with similar ecology. Habitat and diet generalists, cavity nesters and flocking species are more common in urban areas than in other habitats, a phenomenon called biotic homogenization. Yet we do not fully understand why this pattern arises repeatedly: why habitat specialists, or open cup nesters, for example, are at a disadvantage in cities. Citizen science provides an opportunity to first assess the broad patterns in India. Subsequent research across a rural-urban gradient will be key to understanding why and how biotic homogenization occurs.

### Macroecology and Biogeography (9 Questions)

How are birds distributed in space, and what large-scale factors – both historical and current – affect this? The variation in climate and topography across India’s mainland and the two island groups provides an excellent opportunity to answer fundamental questions of this nature, and the answers obtained can help to understand the consequences of global change. New technologies are likely to play an important role in these investigations.

60. What is the geographical variation of rarity and abundance in Indian birds?

**Annotation:** The Indian Subcontinent (including island systems) encompasses a range of geographic and climatic conditions, being at the confluence of three biogeographic realms.

Several species, like the Cinereous Tit *Parus cinereus* and Jungle Myna *Acridotheres fuscus*, can occur in similar habitats across multiple regions in the subcontinent, including southern India, western India, Himalaya, and northeastern India. Why are such species abundant in some geographies but not in others, for example, in southern India but not in western India? What explains the existence of such geographical variation in abundance for some widespread species but not others?

61. What are the drivers of speciation and species range limits in the Indian Subcontinent?

**Annotation:** Speciation is a long-term evolutionary process through which populations evolve to become distinct species. Many drivers of speciation are well known. The Palakkad gap in the Western Ghats is one such example of a geographic barrier that has driven historical speciation. However, many other drivers of speciation are merely suspected, and are poorly delineated and understood. A number of questions emerge from hitherto unexplained species distribution patterns. For example, why does the Eastern Himalaya harbour such a rich assemblage of breeding birds? Is the Narmada river of biogeographic significance? In what ways are drivers of speciation in India now being anthropogenically compromised? Given recent advances in genetic methods (whole genome sequencing) and technology (acoustics), such fascinating questions are calling out to be answered.

62. What are the geographic, climatic and ecological drivers of bird migration?

**Annotation:** Migration is a fascinating annual phenomenon that involves the seasonal movement of birds from one region to another. Bird species vary considerably in the routes they take, when they undertake these journeys, and the habitats they occupy in the different seasons. How do geography, climate (e.g., winds), and ecology interact to determine these routes and maintain them? For example, why do some long-distance migrants to the Western Ghats largely winter only in the northern regions of that mountain range? Do the timing and patterns of monsoon rain and monsoon winds influence migration? How is global change expected to alter spatiotemporal patterns of migration? Many such questions remain unanswered within this critical research theme.

63. What paleo-biogeographic events have given rise to the present-day Indian avifauna?

**Annotation:** As the Indian plate moved away from the Gondwana landmass as an island, it likely had many bird species found nowhere else until it collided with the Laurasian plate about 40 million years ago. What were these endemic species? Are any of these lineages still extant in the subcontinent, or have taxa invading India when it connected with Asia driven them to extinction? How have events such as volcanism or glaciation (ice ages) shaped the diversity and distribution of Indian birds? Studying bird fossils found in various parts of India from diverse paleontological strata (rocks formed at different times in earth’s history) and comparing them to current distributions will help answer this question.

64. What are the ornithological affinities of the Andaman and Nicobar Islands?

**Annotation:** The avifauna of the Andaman and Nicobar Islands appears to bear similarities to Myanmar, Sumatra and the Indian subcontinent. With the emergence of new methods and updated knowledge of bird distributions today, earlier ideas about these affinities need to be reexamined. Phylogenetic studies can help understand the islands’ evolutionary history and highlight their unique bird assemblages *(see also question 73)*.

65. How do species distribution patterns reflect trait-environment relationships?

**Annotation:** The number and types of species vary along environmental gradients such as temperature, precipitation, humidity, and so on. In addition there are variations in traits (such as wing size, bill size, bill type, egg laying pattern, egg size, egg colour, and song) and behaviour both across and within species. For example, the Singing Bushlark *Mirafra javanica* in Africa does not mimic, unlike its Indian counterparts; and Jungle Myna eye colour is golden in its northern populations versus whitish in the peninsula. What is the functional significance of such trait variation, how do traits relate to ecology and environment, and how is this reflected in species distributions?

66. How is global change expected to alter the distributions of common birds?

**Annotation:** Common bird species such as Red-vented Bulbul *Pycnonotus cafer*, Rose-ringed Parakeet *Psittacula krameri*, House Sparrow *Passer domesticus*, Indian Peafowl *Pavo cristatus* and Common Myna *Acridotheres tristis* are highly adaptable species that have widespread distributions across the subcontinent. Global change may favour such generalist species because of the predicted expansion of drier biomes due to various reasons, including land-use and climate change. In the past few decades, for example, Indian Peafowl has expanded its range into the Himalaya and Western Ghats. How do we expect abundances and distributions of common birds to change under these changing circumstances, and what consequences will this have on existing ecological communities?

67. Where are heronries distributed in India and what drives this distribution?

**Annotation:** Heronries harbour important assemblages of wetland birds, providing safe breeding areas for large numbers of individuals. Many heronries are well known as being traditionally protected over centuries by local communities, such as Vedanthangal in Tamil Nadu or Kulik in West Bengal (a large heronry that supports about 40% of the Eurasian Spoonbill *Platalea leucorodia* population). Although many heronries are now protected as Bird Sanctuaries, the locations of these breeding aggregations can also be dynamic rather than static. In the face of both local and global changes, it is critical to know the overall distribution and extent of heronries in India, and why certain habitats and environments are preferred over others. A similar set of questions applies to all breeding aggregations of birds.

68. How do birds respond to climate change and extreme weather events in mountain landscapes?

**Annotation:** Mountain birds, especially those of the Himalaya and Western Ghats, tend to have restricted elevation ranges due to their evolutionary associations with specific climatic envelopes (e.g. temperature and rainfall). Any long-term changes in climatic conditions at specific elevations may make those elevations unsuitable for montane birds by affecting their breeding behaviour, food availability, and demography. In some parts of the Western Ghats, for example, high-altitude grasslands essential for Nilgiri Pipit *Anthus nilghiriensis* are maintained by frost. Any increase in temperature might result in invasion of woody vegetation and therefore the loss of habitat for the species. In the context of similarly sensitive resident and migratory montane species, it is critical to investigate responses (range shifts and local adaptations) to changing climate and higher frequencies of extreme weather events.

### Population Genetics and Evolution (9 Questions)

Understanding the deep history of our birds, and the past causes for current patterns, has been a topic of much work. However, the nature of the questions make them difficult to answer: how can one travel into the past to investigate historical events? Careful observational and experimental studies are now supplemented by modern technologies – including molecular markers and climate-reconstruction models – to address such questions with a greater rigour and historical depth.

69. What are the phylogeographic patterns of disjunctly distributed species across the Indian subcontinent?

**Annotation:** Many species of Indian birds are distributed in a disjunct fashion. For example, the Great Hornbill *Buceros bicornis* occurs in the wet evergreen parts of Western Ghats, and the Himalaya and northeastern India, but is absent from the rest of the Indian Subcontinent. Another curious case is of the *Hypsipetes* bulbuls of the Western Ghats and Sri Lanka, which are sister species to Malagasy *Hypsipetes* and distantly related to Himalayan *Hypsipetes*. Several ideas have been put forward to explain such disjunct distributions, including the Satpura and Brij hypotheses. These hypotheses require robust examination through empirical testing to help us understand historical events that might have influenced the present-day distributions and to predict possible future range contractions.

70. What mechanisms lead to genetic and phenotypic differentiation within a contiguous distribution?

**Annotation:** Physical barriers such as mountains and rivers are known to cause breaks in the continuity of species distributions, with each sub-population subsequently moving along different evolutionary trajectories. This could result in variation in genetic traits or in phenotypic (observable) traits (e.g., physical traits, behaviour, vocalisation) between sub-populations due to adaptation and selection (to specific climatic or environmental conditions), or due to random and neutral processes. In continuously distributed populations, such patterns are diluted because of connectivity. However, they may still arise due to variation in local climates/environments, and other ecological factors such as competition or neutral processes (e.g., geographical distances among sub-populations). Information on such patterns and their causes could help identify barriers/discontinuities in seemingly contiguous populations and improve knowledge of the speciation process.

71. What are the drivers of cryptic speciation in birds?

**Annotation:** Cryptic species are those that outwardly look the same across their range but show genetic, morphological, or behavioural differences between populations that warrant splitting them into two or more species. How do these differences come about? Do behavioural or morphological differences precede genetic differences? Or do gaps (environmental, geographic, or anthropogenic) between populations lead to barriers in gene flow leading to distinct morphology and behaviour evolving in each population over time? Broadly distributed species with high habitat specificity are typically good candidates to study cryptic speciation.

72. What factors drive the evolution of plumage and patterns of moult in Indian birds?

**Annotation:** Birds use their feathers to insulate against the cold, stay dry, fly, attract mates, hide from predators, and communicate with their own and other species. How do all these functions interact to influence the evolution of a species’ plumage? How often do Indian birds moult (replace) their feathers? How does it vary within species or between species? Addressing these questions will require a targeted study of specific species (e.g., pheasants to explore the evolution of ornamental feathers) alongside extensive, large-scale, and long-term bird ringing and monitoring programs, which are essential for understanding variations in moult across the subcontinent.

73. What are the drivers of speciation in the Andaman and Nicobar Islands?

**Annotation:** Island groups tend to be crucibles of evolutionary diversification. As the sea level rises and dips alongside warming and cooling cycles of the earth, the islands lose and gain connectivity amongst themselves and the neighbouring continental landmasses. This results in multiple colonisations, exchange of avifauna and subsequent isolation, potentially resulting in speciation. The avifauna of the Andaman and Nicobar archipelago has been shaped by multiple colonisation events from continental South East Asia (to the Andamans) and the Greater Sunda Islands (to the Nicobars). However, there are oddities – some key groups are particularly abundant, while others are largely absent. A large-scale molecular study in understanding the connectivity between islands and avian speciation events here is a research subject under the larger theme of island biodiversity *(see also question 64)*.

74. What are the phylogenetic patterns in the evolution of female song?

**Annotation:** Traditionally, birdsong has been thought of as a male trait. It has been proposed that this view emerged due to a bias in studies on bird song being conducted disproportionately on species from northern temperate regions of the world. This notion has recently been reversed by evidence emerging that female song is an ancestral trait of modern songbirds.

Given this, it is important to understand the prevalence of female song in Indian birds, and what factors may have driven the evolution, maintenance, or loss of female song in Indian bird species *(see also question 23)*.

75. What is the effect of climate change on the morphology of common bird species?

**Annotation:** Temperature and rainfall are known to impact the evolution of physical traits of birds such as plumage colour and body size. Individuals living in cooler climates are on average larger than those of the same species that live in warmer environments. Birds in rainforests are more colourful than those inhabiting desert ecosystems. Given these patterns, climate change is expected to lead to the evolution of altered physical traits of birds. Since climate change can be viewed as a stress factor, it could also affect the symmetry of physical traits (e.g., wing length) through perturbations in the developmental process. Understanding the short-term selective pressures that could lead to long-term trait evolution is a fascinating topic of climate change research.

76. How does habitat fragmentation affect gene flow in birds?

**Annotation:** Although most birds have the ability to fly, fragmented habitats can nonetheless affect their ability to disperse. For some species this may be severe enough to make isolated populations more vulnerable to local extinction as a consequence of small population sizes. Reduced gene flow across separated populations may lead to inbreeding, resulting in accumulation of deleterious mutations. The reverse is possible too; genetic bottlenecks may eliminate existing deleterious mutations from the local genepool. How do these processes play out in Indian birds, and how does this vary across species?

77. How does habitat alteration affect hybridisation events between species?

**Annotation:** For closely related species, species limits are not absolute and some hybridization is observed in nature. Many congeneric parapatric or sympatric species are known to hybridise in the areas of range overlap, and in some cases the hybrids are fertile, with fitness comparable to the parent species. In other cases, hybrids are inviable or infertile. Habitat alteration may bring potential hybridising species into closer proximity, or in other ways affect the frequency of hybridisation events. How does this play out among Indian species, and what are the consequences of such hybridisation in different scenarios? These consequences would be interesting to study across a range of perspectives, from landscape and biotic interactions to genetic, physiological, and other mechanistic levels. For example, in the latter, are these hybrids facilitated or inhibited by mitonuclear compatibility/incompatibility; or by improved/decreased physiological functions?

### Applied / Economic Ornithology (7 Questions)

Birds provide many ecosystem functions and services including direct and indirect benefits like food, seed dispersal, pest control, pollination, biogeochemical cycling, fertiliser, ecosystem engineering, cultural services, eco-tourism and more. In the Indian context, although we may have some idea of the various functions and services provided by birds, for the most part these have not been studied in detail. This is a multi-disciplinary research theme, cutting across many fields, including biology, chemistry, natural history, economics, modelling, socioeconomics, and biogeography.

78. What are the impacts of bird pollination on Indian plants?

**Annotation:** Animal-pollinated flowers can be ordered along a spectrum with respect to birds: at the one extreme, some may not be visited by birds at all, while others (termed ornithophilous plants) may receive considerable bird visitation. There are very few bird-specialist flowers; most ornithophilous plants are visited by multiple species of flower visitors, which may include birds, bats and insects. Given that all flower visitors may not contribute equally to pollination, what is the relative contribution of bird visitors to pollination and seed-set across different types of flowers?

79. What is the role of birds as biological pest control agents in agricultural ecosystems?

**Annotation:** Birds may act as biological pest control agents in various ways, but much more work on the context and magnitude of these effects is needed. For example, it is widely believed that insectivorous birds are beneficial in controlling crop pests. However, in some situations, insectivorous birds may be a mixed blessing, as they tend to be generalists, and therefore may feed on not just pests but also species like parasitoids and predators (e.g., spiders) that themselves control pests. Similarly, research in other countries suggests that raptors can control or deter rodents in farms, but they may simultaneously deter beneficial insectivorous birds. If birds do indeed control pests, then under what conditions does this occur: which crops are such effects seen in, and does this happen only under certain conditions of habitat matrix (e.g. surrounded by natural habitats) or management practices (e.g. availability of perches)? In addition to conducting broad-scale comparative studies, targeted research is also needed to quantify the impact of a particular predatory species against a specific pest in a specific crop. Results of this kind of research would be of broad applicability in integrated pest management and in conservation.

80. How do colonial breeding waterbirds influence agricultural productivity?

**Annotation:** Colonial breeding waterbirds tend to function as vectors of nutrients, carrying them from feeding areas to breeding and roosting aggregations. Studies elsewhere in the world have measured the quantity of nutrients (particularly phosphorus and nitrogen) in faeces and waterbodies, calculating the annual import of nutrients into adjacent areas by colonial breeding waterbirds, as well as how this affects the water and sediment quality. We know little about these processes in India: in what contexts they occur and to what magnitude. Investigating the role of colonial nesting waterbirds in moving nutrients around, thereby affecting agricultural productivity, adds a further dimension to understanding the ecosystem services birds provide.

81. What management practices can enhance services and manage disservices by birds in agroecosystems?

**Annotation:** Different species of birds (and perhaps even the same species) might provide both positive services (like pollination and pest control) as well as disservices (like crop damage).

Farm management strategies like cropping patterns, retaining semi-natural habitat in and around fields, and artificial or natural perches can help attract certain species by providing suitable conditions for birds. On the other hand, various visual, auditory, tactile, exclusion and olfactory deterrents are used to keep harmful birds away. How can farmers manage the tradeoff between these opposing intentions such that benefits are maximised? This will need to be tested observationally and experimentally in a number of site- and context-specific studies.

82. How does habitat fragmentation affect the ecosystem functions of birds?

**Annotation:** Ecosystem processes are critical for maintenance of biodiversity and for providing services to humans. Birds provide key ecosystem functions, like pollination and seed dispersal, but it is not clear how resilient these functions are in the face of anthropogenic change. With land-use change and habitat fragmentation being key threats to natural ecosystems, it is vital to understand how these processes affect the ecosystem functions fulfilled by birds.

83. What is the magnitude of human-bird conflict, and how can this be minimised?

**Annotation:** Human-bird conflict significantly affects both bird conservation and human wellbeing. Some possible sources of such conflict are birds damaging crops, reducing fish catch, acting as vectors of disease, roosting in large numbers in human-dominated habitats, and colliding with aircraft (see following question). Evaluating these conflicts, assessing their magnitude, and finding out whether indeed they result in perceptible problems requires detailed understanding from different social, cultural, economic, and political perspectives. Global reviews suggest that human-bird conflict is increasing, is acute in developing countries, and is particularly caused by generalist species with growing populations. Studies are needed to understand what bird species are involved in such conflict, and to what extent, and what options are effective to manage these problems.

84. What are the circumstances under which birds collide with aircraft, and what mitigation measures are effective?

**Annotation:** Bird collisions with civilian and military aircraft are a global problem. In India, the Directorate General of Civil Aviation (DGCA) issued a circular in August 2022 titled “Management of potential wildlife hazards at licensed aerodromes”. This circular acknowledges the severity of the problem and highlights the urgency to deal with the hazard. Research is needed to understand how flight schedules and type of aircraft intersect with species size, behaviour, and density to affect collision risk, and how this is affected by the features of the surrounding landscape. Research is needed to suggest suitable management options in different situations, and then to monitor and evaluate their efficacy once implemented. With more aircraft in the skies every year, this is a vital direction for investigation.

### Conservation (17 Questions)

In a rapidly changing world, an important goal is to understand how anthropogenic activities are affecting birds and what can be done to minimize or reverse these effects. Research is a vital component of this endeavour because identifying threats, finding and testing solutions, and monitoring the outcomes of interventions all require rigorous science. Conservation, which involves protecting habitats, species, and ecological processes, is most effective when guided by research that directly addresses a problem and yields actionable results. By integrating scientific findings with conservation efforts, we can develop targeted strategies to protect bird populations and the ecosystems they depend on.

85. How do large-scale green energy projects affect birds?

**Annotation:** Climate change mitigation projects often involve setting up large-scale green energy projects, such as wind and solar power farms in supposedly barren areas and wastelands. Many such sites are actually grasslands or other Open Natural Ecosystems (ONEs) that are degraded or destroyed in the process. Apart from the power farms themselves, the accompanying infrastructure (like, roads and power lines) can cause further problems. For example, in Rajasthan, large-scale wind energy turbines and power transmission lines are implicated in elevated mortality rates of the Critically Endangered Great Indian Bustard. There is only a rudimentary understanding of the exact type and magnitude of these sorts of projects on birds, and much remains to be done.

86. What is the impact of free-ranging dogs and cats on birds?

**Annotation:** The impacts of free-ranging dogs and cats on birds and other wildlife appear to be increasing over time, but in India the matter has not received the research attention it deserves. Consequently, we have no rigorous estimates of the magnitude of the problem (e.g., number of birds killed), nor what might underlie variation in these impacts, let alone what options are available for effective mitigation in different types of habitats including cities, towns, villages and agricultural landscapes. The answers to these questions are needed if India is to devise sensible policies related to free-ranging dogs and cats, which take into account conservation needs as well.

87. How do large-scale commercial fishing activities affect shore and pelagic birds?

**Annotation:** Large-scale mechanised fishing, aided by satellite-based observations that provide information on fish aggregation sites, together with advanced tools and gear are depleting marine fisheries at the global level. Some of these activities are illegal, unreported and unregulated. They are likely to be adversely affecting oceanic and pelagic birds, through both depletion of food and inadvertent mortality from fishing gear (nets and hooks). The status of pelagic birds in Indian waters is poorly known, and effects of fishing may already be driving populations down unnoticed, so there is considerable urgency to tackle this question.

88. How does linear infrastructure development impact birds?

**Annotation:** Linear infrastructure, such as power transmission lines, multilane highways and expressways, rapid railway networks, large canal systems and interlinking of rivers impact biodiversity not only through loss and fragmentation of habitats, but also by creating barriers to the movement of birds and animals. In the long term, infrastructure like roads may be ‘contagious’, altering land use in more and more natural habitats. The various short- and long-term impacts of such infrastructure (including collisions with vehicles and powerlines, and restrictions on movement) need to be properly understood if we are to make sensible decisions about planning and mitigation.

89. What are the impacts of plastic waste on waterbirds?

**Annotation:** Plastic waste is common everywhere in waterbodies, with possibly severe impacts on many organisms. Macroplastics in the water are consumed directly by birds that feed on phytoplankton and zooplankton. Microplastics also enter the food chain and can bioaccumulate in fish, which are then consumed by a large variety of birds. Birds get entangled in plastic rings, bottles, and other products, often starving or drowning in the process. Discarded fishing gear (‘ghost nets’) is especially lethal. Millions of fishing lines with hooks litter our oceans, a hazard for pelagic birds and shorebirds. Research has lagged behind in understanding the magnitude of this large and growing problem, and in finding appropriate solutions.

90. How does hunting and illegal trade affect birds in India?

**Annotation:** Illegal wildlife trade is a serious crime with a global turnover of billions of dollars annually. Commercial, recreational and consumption-based hunting is prevalent in many parts of India despite its legal prohibition by the Indian WildLife (Protection) Amendment Act 2022. The existing work on this issue has mostly documented species (and numbers) in a subset of markets, or has collated information on seizures of illegal consignments of birds. The cumulative numbers derived in this way are clearly an underestimate of the true magnitude of offtake from the wild. Still, it is possible that many wild species are common enough and breed fast enough to withstand considerable offtake. Given this, we need detailed studies that estimate the overall effect of hunting and trade on the populations of various species, on hotspots of hunting and trade, and on species that are particularly targeted. These will allow better assessment of the magnitude of conservation concern, and assign priorities for action accordingly.

91. How do changes in carcass disposal practices affect the distribution and abundance of raptors in India?

**Annotation:** Traditionally, carcasses of cows or other livestock have been disposed of in the open, to be fed upon by natural scavengers. This practice appears to have reduced in recent years, with carcasses increasingly being buried rather than discarded in the open, possibly in response to decreasing vulture populations. What are the consequences of these changes for vultures and other scavenging raptors? This group of birds has experienced particularly heavy decline in the past couple of decades, and any recovery may well be dependent on the availability of food.

92. What are the effects of tourism on birds and their habitats?

**Annotation:** Tourism in India is increasingly linked to nature. While this is a welcome development, rising tourism demands better infrastructure (e.g., roads, hotels, electricity), and resources (e.g., water and fuel, including fuelwood). These demands increase the pressure on natural habitats, potentially affecting bird populations. Vehicles and people can generate disturbances that affect the behaviour of birds. Photography and associated tools (e.g., song playback, hides, feeders) may have additional effects. How all these effects act individually and in combination to impact bird behaviour, food availability, breeding, and populations is a topic of urgent research.

93. What are the effects of audio playback on birds in India?

**Annotation:** Birds use song and calls to communicate with each other, and playback of recorded audio is commonly used in ecotourism to bring birds into the open for better views by birdwatchers and photographers. Playback is also used in research to investigate the function of different sounds, and as an assay in population monitoring of pheasants and owls. Playback for ecotourism has been criticised, but the actual short- and long-term impacts on behaviour and population dynamics are poorly understood. This calls for careful research on estimating the magnitude and context of possible negative effects.

94. How do cultural traditions and people’s attitudes affect birds in India?

**Annotation:** The rich cultural diversity of India is reflected in its large number of festivals and cultural activities. Some of the accompanying practices are known to harm birds: these include setting off fireworks and flying kites with glass *manja*. Other activities, for example the deployment of decorative lighting on a large scale, may possibly cause harm (through increased light pollution), but this is not well understood. Feeding birds (pigeons and crows particularly) is widespread, but again the impact on bird communities as a whole is not clearly understood.

Specific taxa may be harmed by both negative (e.g., owls) as well as positive (e.g., Indian Rollers *Coracias benghalensis*) beliefs of communities. Finally there may be some cultural practices and attitudes that promote conservation (e.g., for species like peafowl, sparrows, and cranes). These varied attitudes and practices are important to investigate deeply, for a fuller understanding of the conservation implications of various cultural beliefs, thereby pointing the way to meaningful societal engagement.

95. What indigenous knowledge exists about birds, and how can it contribute to conservation?

**Annotation:** Indigenous knowledge about birds is encapsulated in various forms that are passed down the generations, including art, song and folklore. Documenting and compiling this knowledge ensures that it can pass to future generations of the same community as well as the larger citizenry. Indigenous knowledge can be used as a springboard for better understanding the natural history of various species as well as to generate ideas for how to more meaningfully understand human-nature relationships, and their implications for conservation.

96. What impact have introduced or escaped exotic birds had on other birds in India?

**Annotation:** From a global perspective, introduced or escaped exotic birds can cause substantial damage and threaten native biodiversity in various ways. However our understanding of this in the Indian context is minimal. A number of bird species have been introduced from mainland India to the Andaman and Nicobar Islands, where their impact on native and endemic species needs to be assessed. On the mainland, some species (e.g., Alexandrine Parakeet *Psittacula eupatria*) appear to have been introduced into new regions from elsewhere in India – this gives an opportunity to study what might help versus hinder establishment in new areas.

97. Are artificial nest boxes effective in enhancing the survival rates of target species?

**Annotation:** There is growing global concern regarding the loss of nesting spaces for birds that nest in trees and other cavities. In India, over 200 species are known to use tree cavities.

Providing artificial nest boxes can be an effective means to improve the population status of such species, especially in agriculture and urban landscapes, and a number of efforts are underway to make and install nest-boxes across India. However, the long-term efficacy of artificial nest boxes in improving nesting rates and outcomes, and thereby the population status of species, is poorly understood and requires research attention.

98. What are the causes of mass mortality of birds?

**Annotation:** The frequency and magnitude of incidents of mass mortality of birds are increasing globally and in India as well. The causes behind these events are varied, including disease, biotoxicity, acute poisoning, sudden changes in weather, food shortage, wildfires, linear intrusions, and more. There is an urgent need for the systematic documentation of such events, as well as detailed investigation of underlying reasons and potential prevention.

99. How should bird species be prioritised for conservation in India?

**Annotation:** Over 1210 regularly occurring bird species have been reported from India, but clearly these are not all of equal conservation interest. On what basis should conservation priority be assigned to them? There may be a number of considerations, including risk of extinction (possibly indexed by population size and trend, range size and trend, and magnitude of known threats), endemicity, and functional role (e.g., as pest control agents, or seed dispersers). Migratory species may require special attention, particularly those for whom the Indian subcontinent comprises all or a large fraction of their non-breeding range. Common (abundant and widespread) species typically do not receive conservation attention, but their functional importance is likely to be higher than rare or threatened species. There are many gaps in the information underlying conservation prioritisation, and these require dedicated research attention.

100. How effective are various conservation measures in conserving Indian birds?

**Annotation:** Several conservation measures exist to protect birds in India, including the designation of special areas (Protected Areas), laws to protect specific species and habitats

(e.g. The Wild Life (Protection) Act [WLPA], the Forest Conservation Act, Environment Protection Act, Coastal Regulation Zone among others), and targeted conservation interventions for specific species (including bustards and vultures). Other areas are designated as being particularly rich in birdlife or important for conservation, but such a designation does not confer legal protection. These include Important Bird and Biodiversity Areas (which can be within existing PAs or outside), World Heritage Sites (important cultural, historical and conservation areas) and Ramsar Sites (important wetland areas). India is also a signatory to several international conventions that protect birds (CITES, Convention of Migratory Species, among others). The WLPA also designates species and groups as particularly protected (in its Schedules). At a broad scale, how effective have these various policy and conservation interventions been? A critical assessment of the efficacy of these measures, and identification of opportunities beyond them, is called for.

101. How do policies unrelated to conservation affect birds?

**Annotation:** Some of India’s policies are focused on biodiversity conservation, such as the designation of Protected Areas. But there are a number of policies that are motivated by other goals, yet they may affect conservation profoundly. For example, the Integrated Wasteland Development Programme appears to designate certain critical ecosystems as ‘wastelands’, based on their perceived lack of value of these natural habitats to human livelihoods and economies. Various policies around rivers and river interlinking, port construction, renewable energy, and agrochemicals have direct and indirect consequences for conservation, as have decisions taken by land-holding administrative departments and Ministries (e.g. Railways, Agriculture, Revenue, Defense, Industry). The implications of these policies and decisions need to be studied if we are to have a broad understanding of how bird conservation is affected by larger social and political forces.

## Discussion

A group of 39 individuals collectively curated 101 key research questions in the field of ornithology in India with contributions from 309 people across careers and career stages. The final list of questions was organized into 9 thematic areas. While the number of questions per theme varies, this distribution does not imply a hierarchy of importance. Together, these questions provide a concise, community-driven framework that captures both the breadth and interconnectedness of ornithological research in India.

### Overarching themes

Although the questions were broadly grouped into nine subject areas, a few themes emerged repeatedly across all subject areas, cutting across disciplinary boundaries and reflecting shared priorities within the community.

In particular, a number of questions revolve around the effect of human activity on the physiology, behaviour and population ecology of birds. Such questions found their way into multiple subject categories, and relate to effects of light, noise and air pollution; land use change, including infrastructure development; agricultural intensification; and changes in hydrology.

Among these, direct human–bird interactions emerged as the most frequently assigned theme across the 101 selected questions (Figure 2). These include questions on public health (e.g., birds as carriers of antibiotic resistance), ecosystem services (e.g., bird pollination), and infrastructure challenges (e.g., bird strikes at airports). Several questions also relate to socio-cultural dimensions of bird–human relationships, such as traditional hunting practices, nest box provisioning, and the role of indigenous ecological knowledge. The diversity of topics under this theme reflects the growing recognition that birds and humans are deeply intertwined across ecological, economic, and cultural contexts.

**Fig 2:**
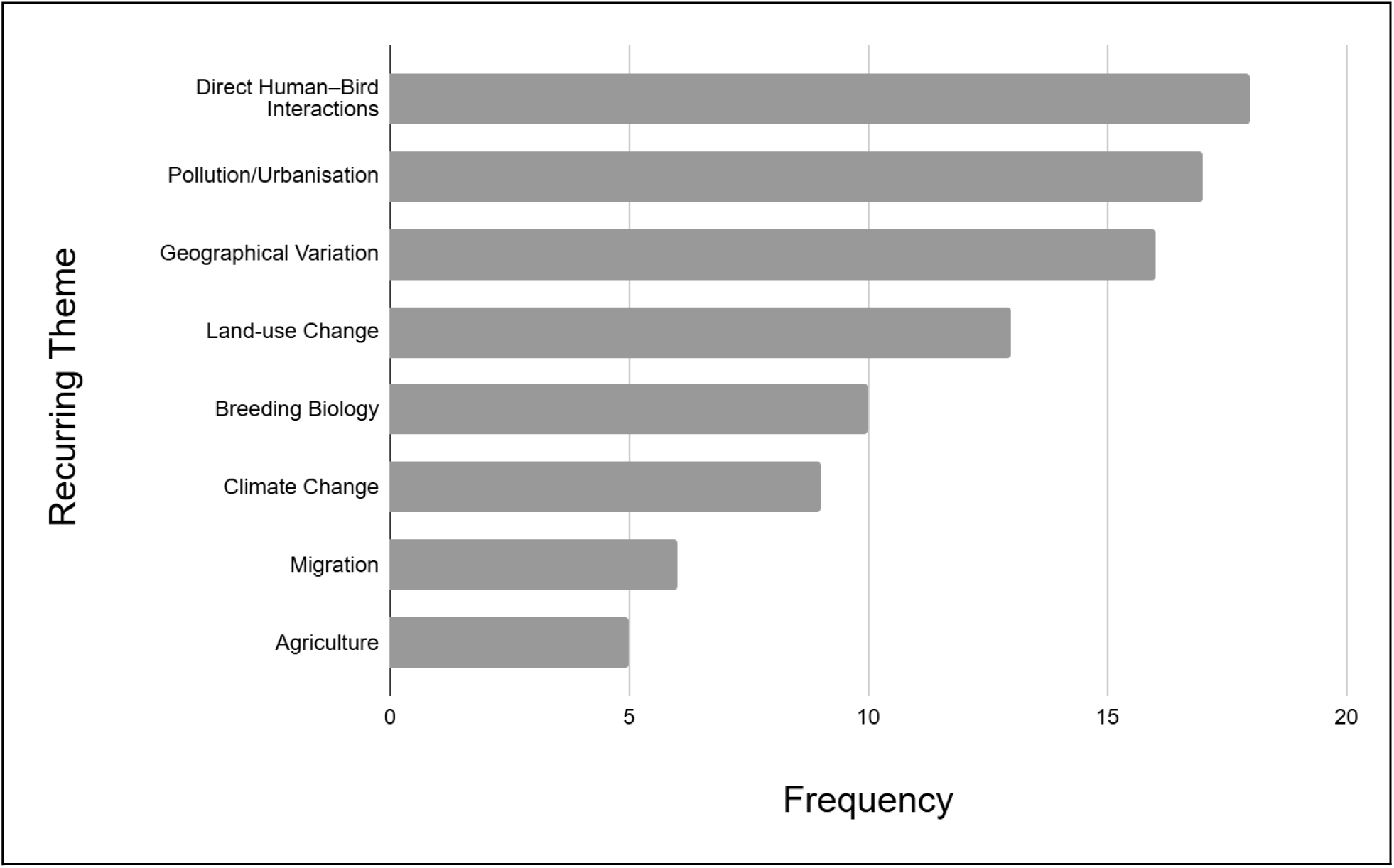
Representation and recurrence of overarching themes across the 101 shortlisted research questions. The figure depicts broad themes that emerged repeatedly across questions from different subject areas. Of the 101 questions, 11 were assigned more than one theme, and such cases are counted separately in the graph above. For example, a question examining how climate change affects bird migration routes and land-use interactions was assigned to both Climate Change and Land-use Change.

In addition to direct human–bird interactions, there were other recurring themes (Figure 2). A substantial number of questions focus on pollution and urbanisation, including the effects of light, air, and noise pollution on bird behaviour and physiology, and the broader ecological impacts of expanding urban infrastructure. Questions related to geographical variation span multiple disciplines—from population differentiation and trait variation to species distributions—highlighting the importance of understanding spatial heterogeneity in bird ecology and evolution across the Indian subcontinent. Land-use change is another prominent theme, with questions examining how agriculture, plantations, dams, mining, and other forms of land transformation influence bird communities, habitats, and movement. A significant subset of questions also deals with breeding biology, addressing both proximate factors such as cues for breeding and nesting success, as well as broader evolutionary and ecological patterns. Finally, many questions explore the impact of climate change, including how warming and shifting weather patterns affect migration, morphology, reproduction, and habitat availability—along with concerns about the ecological effects of climate mitigation measures like renewable energy infrastructure.

Broadly speaking, our list contains a large fraction of questions that have a conservation or other applied motivation compared with those that might be considered more ‘basic’ or ‘fundamental’. For example, the list contains relatively few questions in Physiology and Disease Ecology or in Population Genetics and Evolution. The skew in subject matter is unintentional, and we believe it is unlikely to be solely a result of the composition of the contributors. Topics that are specialised in nature, or that require complex techniques and expensive equipment are understandably underrepresented in the larger landscape of ornithology research in India, and in our list of questions as well. This can be contrasted with observation-based field studies of behaviour and community ecology, which appear to be more frequent. We emphasise that although mechanistic studies in physiology, population genetics, etc, may be difficult to carry out, they are nonetheless crucial to answer both basic questions about bird biology as well as to create a foundation to tackle applied problems arising from, for example, climate and land-use change.

### New methods and approaches

It is worth pointing out that new technologies, tools, methods, and approaches are increasingly being used in Indian ornithological research. Field-based observations are being supplemented by recent methods developed for both lab and field. For example, there is a growing use of capture-based methods to study demography; of genetic tools to understanding speciation, species delimitation, and phylogeography; of fossil studies to investigate historical ranges; and of molecular techniques to study avian physiology and disease. Tagging and telemetry-based movement studies are also becoming increasingly popular; these require funds, permits, and trained personnel. As in other parts of the world, ornithology in India is evolving to be more interdisciplinary and collaborative, ensuring that the most suitable tools and methods can be brought in to address the question at hand.

Another approach that is gaining traction in India is where enthusiasts and professional scientists work together towards a common research goal in what is known as ‘citizen science’. Although citizen science is not new in India or elsewhere, it appears to be in a phase of rapid growth. This approach has resulted in the generation of data on bird populations and distributions at a spatial and temporal scale previously unimaginable. On public platforms like eBird, millions of observations are contributed from across the country every year. These data have been used as the basis of periodic conservation assessments like the State of India’s Birds reports (e.g., SoIB 2023), but the massive database is still underutilized in addressing a plethora of topics and questions about distributions, communities, migration and more. Just as importantly, the ideas and technologies enabling citizen science have stimulated groups of like-minded enthusiasts to design and implement their own projects, for example, the series of bird atlas studies across India (e.g., Praveen and Nameer 2021). The involvement of non-professionals in Indian ornithology is demonstrated by the fact that as many as 48% of respondents to the form circulated for this study described themselves as Nature Enthusiasts.

### Enabling Indian ornithology

Looking ahead, the continued growth of ornithological research in India will depend on the convergence of several factors across different sectors.

First, enabling a vibrant research environment within and across scientific institutions requires consistent and reliable funding. This is crucial for developing and maintaining research field stations, laboratories and maintenance of equipment. While financial support is scarce overall, funds for field infrastructure and work are particularly hard to raise.

Second, regulatory frameworks and policies have a key role to play in the research environment in the country. Policy can determine the kind of research possible and can severely limit or greatly boost research. We can identify specific obstacles in the form of the difficulties in accessing study sites as well as study species due to restrictions on access specified in the WildLife (Protection) Amendment Act 2022. In particular, permits to conduct research in protected areas, or to capture birds are hard to come by, and much vital research is therefore impossible to conduct. Despite this being a long-recognised barrier to wildlife research in India (Madhusudan et al. 2006), there has been no discussion or resolution among the key stakeholders - researchers, policymakers, wildlife managers, funders, and others.

Third, collaborations enable interdisciplinary research and bring together different ideas and diversity that can shape the research and education for the future of ornithology in India.

Building a collaboration-friendly environment as a community will benefit the integration of diverse disciplines to further improve the quality and quantity of ornithology research in India.

Fourth and finally, the value of citizen science must be recognised by both researchers as well as science funders, administrators and policymakers. The potential and promise of citizen science is not only as a way of scaling traditional approaches to research but, vitally, of also reimagining new ways in which science and research can be led and carried out by the people at large – and in this form, has the potential to strengthen and transform Indian ornithology.

## Conclusion

In this collective effort, we have curated a list of research questions in Indian ornithology that we suggest are of considerable interest and importance. In doing so, we have drawn from a variety of audiences, from enthusiasts and researchers, to students, managers and policymakers. The resulting list of questions necessarily reflects the understanding and preferences of those who contributed questions as well as those who curated them (the authors of this paper). We do not claim that these are the only vital questions in Indian ornithology, yet at the same time we feel they deserve considerable time and attention. We hope these questions, along with the annotations, will inspire and stimulate new project ideas for students, researchers, and citizen scientists, while also guiding funders, managers, and policymakers in identifying research areas that need support and encouragement.

## Supporting information

Questions in the Google form (Open call to recommend questions)

## Acknowledgments

We are grateful to those who contributed questions to this project; all contributors are listed in the supplementary material to this paper. We thank them all for their thoughtful contributions and the collaborative spirit that made this project possible. We are grateful to Uma Ramakrishnan, the National Centre for Biological Sciences, and the Institute for Stem Cell Science and Regenerative Medicine for their help with the in-person meeting held as part of this project.

Travel and accommodation was supported by the Nature Conservation Foundation.

## Author Contributions

Initiating group: Rajah Jayapal, Manjari Jain, Farah Ishtiaq, Devica Ranade, Suhel Quader

Subject Groups:

Applied/Economic ornithology: Govindan Veeraswami Gopi*, Hilloljyoti Singha, Rajah Jayapal, Subbu Subramanya

Behaviour: Priti Bangal*, Anil Kumar, Dhanashree Paranjpe, Gopinathan Maheswaran, Manjari Jain, Sahas Barve, Suhel Quader

Community Ecology: Rohit Naniwadekar*, Kulbhushansingh Suryawanshi, Monica Kaushik,

Mousumi Ghosh, Prachi Mehta, Raman Kumar

Conservation: Ajai Saxena*, Malyasri Bhattacharya*, P Jeganathan, Parveen Shaikh, Peroth Balakrishnan, Shomita Mukherjee, Suresh Kumar

Habitat ecology: Girish Jathar*, Aparajita Datta, Ankita Sinha, Bhoj Kumar Acharya, Malvika Onial, Monica Kaushik, PO Nameer

Macroecology and Biogeography: Ashwin Viswanathan*, Ajai Saxena, Ankita Sinha, Bhoj Kumar Acharya, Madhumita Panigrahi, Vivek Ramachandran

Natural History, Systematics and Evolution: Ashish Jha*, Anil Kumar, Balakrishnan Peroth, P Jeganathan, Praveen J, Sahas Barve, Shomita Mukherjee, Suresh Kumar

Physiology & Disease Ecology: Anusha Shankar*, Farah Ishtiaq, Manjari Jain, Soumya Iyengar

Population Ecology: Parveen Shaikh*, Govindan Veeraswami Gopi, Girish Jathar, Malyasri Bhattacharya, Rajah Jayapal

Working Groups

● Metadata and Workflow: Ankita Sinha, Anusha Shankar, Priti Bangal
● Scoring Guidelines: Aparajita Datta, Monica Kaushik, Mousumi Ghosh

Manuscript Preparation

● The manuscript was written and revised by Anusha Shankar, Devica Ranade, Manjari Jain, Priti Bangal, Rajah Jayapal, Rohit Naniwadekar, Suhel Quader
● All authors reviewed and provided feedback on the final manuscript.

*subject group moderators

