## Supplementary material for "What’s next for Indian Ornithology? 101 key research questions": Questions in the Google form (Open call to recommend questions)

The form which was circulated among the larger audience included the following questions:

1. Name
2. Profession/ Your involvement with ornithology (Select one)
    - Ornithology/Ecology student
    - Ornithology/Ecology researcher or teacher (post PhD)
    - Wildlife/Environment manager or policy maker
    - Birdwatcher
    - Other nature enthusiast (eg, wildlife photographer)
    - Other

3. Please enter your suggested research question here

4. Broad subject under which the above research question falls (Select one)
    - Natural History
    - Systematics and Evolution
    - Macroecology and Biogeography
    - Habitat Ecology
    - Community Ecology
    - Population Ecology
    - Behaviour
    - Physiology
    - Disease Ecology
    - Conservation
    - Applied/Economic Ornithology

5. Apart from the subject area you have chosen above, please add up to 5 keywords of your choice
